# Signatures of the evolution of parthenogenesis and cryptobiosis in panagrolaimid nematodes

**DOI:** 10.1101/159152

**Authors:** Philipp H. Schiffer, Etienne G.J. Danchin, Ann M. Burnell, Anne-Marike Schiffer, Christopher J. Creevey, Simon Wong, Ilona Dix, Georgina O’Mahony, Bridget A. Culleton, Corinne Rancurel, Gary Stier, Elizabeth A. Martínez-Salazar, Aleksandra Marconi, Urmi Trivedi, Michael Kroiher, Michael A.S. Thorne, Einhard Schierenberg, Thomas Wiehe, Mark Blaxter

## Abstract

Most animal species reproduce sexually, but parthenogenesis, asexual reproduction of various forms, has arisen repeatedly. Parthenogenetic lineages are usually short lived in evolution; though in some environments parthenogenesis may be advantageous, avoiding the cost of sex. *Panagrolaimus* nematodes have colonised environments ranging from arid deserts to arctic and antarctic biomes. Many are parthenogenetic, and most have cryptobiotic abilities, being able to survive repeated complete desiccation and freezing. It is not clear which genomic and molecular mechanisms led to the successful establishment of parthenogenesis and the evolution of cryptobiosis in animals in general. At the same time, model systems to study these traits in the laboratory are missing.

We compared the genomes and transcriptomes of parthenogenetic and sexual *Panagrolaimus* able to survive crybtobiosis, as well as a non-cryptobiotic *Propanogrolaimus* species, to identify systems that contribute to these striking abilities. The parthenogens are most probably tripoids originating from hybridisation (allopolyploids). We identified genomic singularities like expansion of gene families, and selection on genes that could be linked to the adaptation to cryptobiosis. All *Panagrolaimus* have acquired genes through horizontal transfer, some of which are likely to contribute to cryptobiosis. Many genes acting in *C. elegans* reproduction and development were absent in distant nematode species (including the Panagrolaimids), suggesting molecular pathways cannot directly be transferred from the model system.

The easily cultured *Panagrolaimus* nematodes offer a system to study developmental diversity in Nematoda, the molecular evolution of parthenogens, the effects of triploidy on genomes stability, and the origin and biology of cryptobiosis.

## Introduction

Despite the general prevalence of sexual reproduction in animals, parthenogenesis has evolved in many taxa. Parthenogenesis, or, more strictly the cessation of outcrossing after meiotic recombination, is thought to result in the gradual accumulation of deleterious mutations through Muller’s ratchet which (Lynch et al. 1993). This would condemn non-recombining parthenogens to long-term evolutionary failure. However, several observations argue against such a simple view: parthenogenesis is a phylogenetically persistent trait in the tree of animals. It must have evolved independently multiple times as a strategy of reproduction. Some asexual taxa (Felsenstein 1974), such as the bdelloid rotifers or darwinulid ostracods, show a surprisingly high rate of speciation and long-term survival without sexual reproduction. The recruitment of recombination-like mechanisms such as gene conversion and inter-individual horizontal gene transfers (Signorovitch et al. 2015; Debortoli et al. 2016) may have rendered Muller’s ratchet ineffective. Furthermore, many parthenogens have been observed to amplify the number of gene or genome copies through allo- or autopolyploidy, and this may buffer lineages against mutation accumulation in essential genes (Archetti 2004). Comparisons of related sexual and asexual taxa are required to understand the origins and persistence of asexuality, and the effects of asexuality on genome evolution.

Asexual reproduction has frequently been associated with organisms found in extreme or peripheral environments, such as arid deserts where water availability is ephemeral and unpredictable or polar ecosystems where much of the annual cycle involves temperatures well below zero °C. Parthenogens have been shown to colonise ephemeral or widely spaced niches as single reproductive propagules (Lynch 1984). In turn, organisms that inhabit or colonise extreme environments, are often cryptobionts. Cryptobiosis is the ability of an organism to survive with no evidence of active metabolism under conditions inimical to life. Cryobionts survive freezing, avoiding lethal ice-crystal disruption of biological membranes, while anhydrobionts survive loss of effectively all their body water. Cryobionts are frequently also anhydrobiotic, as ice sublimation in very low temperatures can freeze-dry. The mechanistic bases of cryptobiosis have been explored in some species (*Aphelenchus* nematodes (Reardon et al. 2010), the chironomid *Polypedilum vanderplanki* (Gusev et al. 2014), and the tardigrade *Ramazzottius varieornatus* and *Hybsibius dujardini* (Hashimoto et al. 2016)) and individual molecular players identified. Intrinsically unstructured proteins and trehalose are important molecular components in anhydrobiotic mechanisms. Intrinsically disordered proteins do not adopt a stable secondary structure in solution, and are thought to replace structural water on dehydration. They include the late embryogenesis abundant (LEA)-like proteins (Pouchkina-Stantcheva et al. 2007; Tyson et al. 2012; Gusev et al. 2014), and the tardigrade-specific SAHS, MAHS and CAHS family proteins (Hashimoto et al. 2016). Trehalose is likely to act in a similar way. While some species are able to undergo cryptobiosis rapidly at any stage of development (Hashimoto et al. 2016), others require extensive preconditioning, presumably to activate physiological systems that produce anhydro- or cryo-protectants.

Both asexuality and cryptobiosis are advantageous to organisms colonising extreme habitats. However, it is currently unclear whether they are mechanistically unlinked or whether the mechanisms giving rise to asexuality may have made acquisition of cryptobiosis traits easier. In the bdelloid rotifers it has been suggested that DNA breaks resulting from the stress of cryptobiosis coupled with gene-conversion may have promoted the acquisition of foreign DNA (horizontal gene transfer), which in turn conferred improved cryptobiotic potential (Debortoli et al. 2016).

Many asexual and/or cryptobiotic taxa of nematodes have been described. The Panagrolaimidae include both sexual (amphimictic) and parthenogenetic taxa. Many, perhaps all, panagrolaims are cryptobionts, with both cryobiotic and anhydrobiotic abilities, and are found in extreme environments in polar, but also in temperate regions (Shannon et al. 2005). Although parthenogenetic species are widely distributed around the globe, a single origin of parthenogenesis in *Panagrolaimus* is likely (Lewis et al. 2009). Still, their exact evolutionary age is unknown. *Panagrolaimus* species are very good cryptobionts: some species can survive very rapid desiccation, and many species can survive desiccation at any life- stage without preconditioning (Shannon et al. 2005).

Here, we compared the genomes and transcriptomes of parthenogenetic and amphimictic panagrolaims, including a *Propanagrolaimus* species which is not a good cryptobiont. We describe a single, recent origin of asexual *Panagrolaimus* species, possibly involving polyploidisation, and estimate the previously unknown age of parthenogenesis in the genus. We identified candidate genes that may be important for cryptobiosis, some of which are likely to have been acquired horizontally. The genome and transcriptome data and analyses presented here constitute a resource for future functional studies of the evolutionary origins of parthenogenesis and cryptobiosis.

## Results

### New genomes and transcriptomes from panagrolaimid nematodes

We sequenced and assembled *de novo* the genomes and transcriptomes of five panagrolaimid nematodes (**Fig. 1**) using contamination- and heterozygosity-aware methodologies. We reassembled the genome of *Panagrolaimus davidi* DAW1, which was highly fragmented and contaminated with a considerable amount of bacterial DNA in the original assembly described in (Thorne et al. 2014).

**Figure 1.**
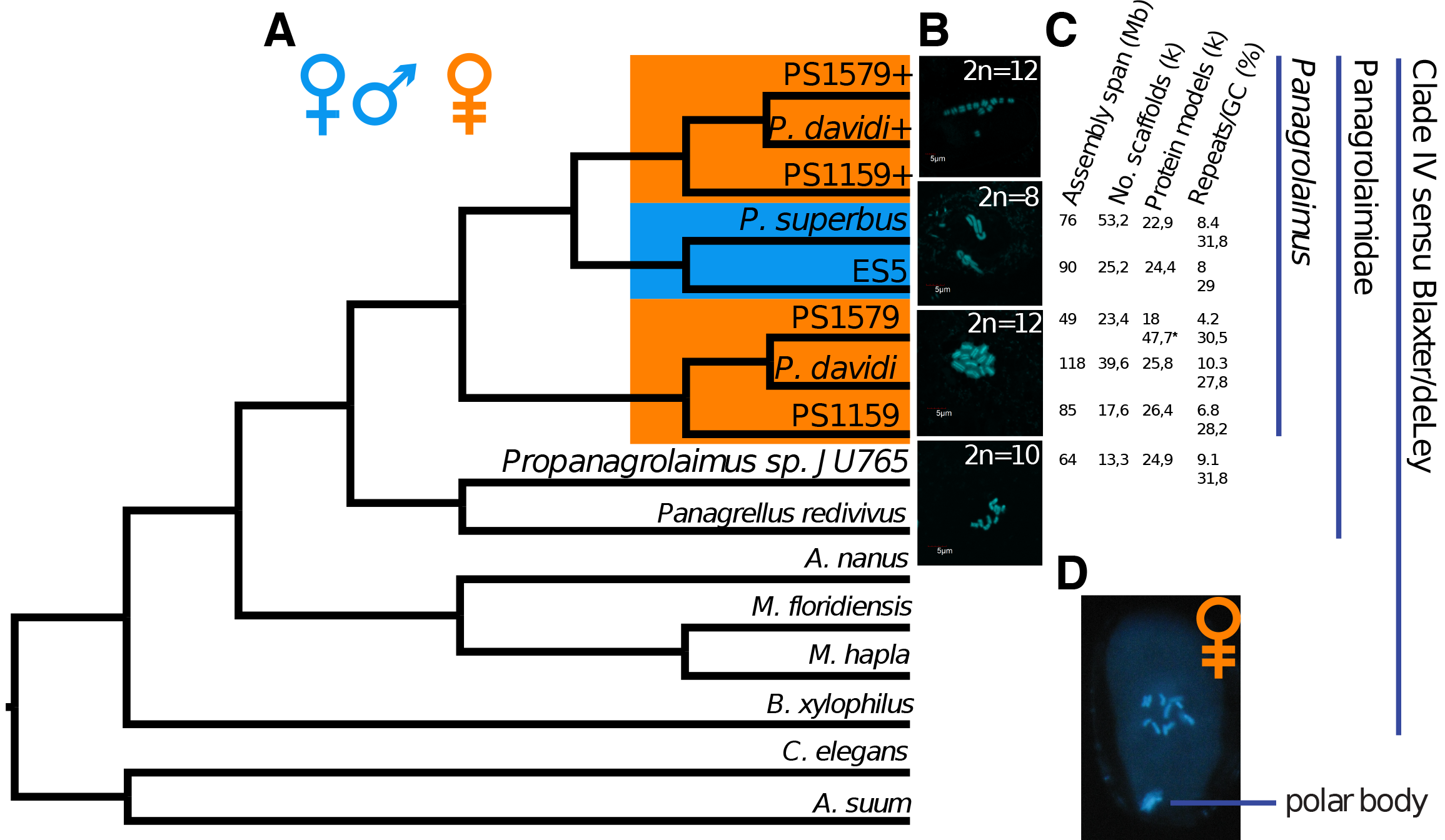
MUL-Tree cladogram, karyotypes, and genome assemblies. **A.** A MUL-trees based gene-tree reconciliation based on 15,000 trees from Orthofinder found the most parsimonious phylogenetic position for proteins where the parthenogenetic species had multiple copies in a given orthologous group to be outside of a clade containing our gonochoristic and parthenogenetic *Panagrolaimus*. This topology supports the assumption of polyploid hybrids. **B.** All analysed parthenogenetic species had 12 chromosomes in comparison to 8 in the diploid state of the amphimictic species. The outgroup *Propanagrolaimus* has 2n=10. **C.** Assembly and gene prediction metrics. P. sp. PS1159 and *P. davidi* both have more predictions than the sexual species. The low coverage PS1579 genome appears to lack regions of the genome, and we thus included transcriptome based gene predictions(*) directly into downstream analysis. **D.** The presence of polar bodies in the parthenogenetic panagrolaims confirms the presence of functional meiosis.

We independently estimated the genome sizes of strains in culture using Feulgen Image Analysis Densitometry (FIAD) and quantitative PCR (RT-D-PCR). For the amphimictic *Panagrolaimus* isolates diploid genome sizes were estimated to be between 0.14 (P. sp. ES5, using FIAD) and 0.18 (*P. superbus*, using RT-D-PCR) pg DNA per nucleus, or ~140-180 Mb. For the parthenogens *(P.* sp. PS1159, PS1579), we obtained FIAD values between 0.23 and 0.27 pg DNA per nucleus (~230-270 Mb), i.e. ~50% larger than the amphimictic strains. The diploid genome size of *Propanagrolaimus* sp. JU765 was estimated at 0.13 pg DNA per nucleus (130 Mb).

The spans of the haploid genome assemblies of *P.* sp. ES5 and *P. superbus* (90 Mb and 76 Mb respectively, **Fig. 1**) are congruent with the sizes estimated by densitometry and PCR. Similarly, the haploid genome span of the hermaphrodite outgroup *Prop.* sp. JU765 was 64 Mb, in line with the FIAD estimate. The correspondence between assembly span and measured nuclear DNA mass was less clear-cut in the parthenogens. The assembly span of *P.* sp. PS1159 was only 85 Mb, similar to that of the haploid genomes of the sexual species but one third of the physical measurement. The assembly span for *P. davidi* was 175 Mb. After cleaning and removal of some duplicate contigs stemming from heterozygosity the final scaffolding measured 118 Mb. Notably, the initial assembly span of 175 Mb matches into the genome size range of ~140-180 Mb measured for the sexual species and is approximately 2/3 the size of the 270 Mb measured for the parthenogenetic species.

We karyotyped 15 *Panagrolaimus* species and found that all the amphimictic species had a diploid set of 8 chromosomes (*2n=8*), while all the parthenogenetic species had 12 chromosomes in total (*xn=12*) (**fig. 1**, also compare to supplementary fig. 1). This could correspond to a shift in haploid chromosome number from *n=4* to *n=6*, or triploidy (where *n=4* is maintained but somatic nuclei are *3n*). Observation of polar bodies in the parthenogenetic species shows that meiosis is present (**fig. 1**).

### Evidence for triploidy in parthenogenetic *Panagrolaimus*

All assembled genomes showed high completeness, comparable to other de-novo sequenced genomes, for the eukaryotic gene set in the BUSCO pipeline (Simao et al. 2015) (see Supplementary Excel file BUSCO results). We performed gene-finding on the new and reassembled genomes, and predicted proteins from the transcriptomes of *P.* sp. DL137 and *P.* sp. PS1579. The number of gene models for the panagrolaimid species had poor correlation with their genome sizes (**fig. 1**). We retrieved domain annotations for 77% - 88% of the predicted coding genes in each species. These were orthology-clustered and analysed with kinfin (Laetsch and Blaxter 2017), including proteomes from *Propangrolaimus* sp. JU765 (this work), and other Tylenchina (Clade IV) nematodes (*Panagrellus redivivus (Srinivasan et al. 2013), Meloidogyne hapla (Opperman et al. 2008), Acrobeloides nanus* (Schiffer et al. 2018) and *Bursaphelenchus xylophilus (Kikuchi et al. 2011)*) as outgroups. In a polyploid system extra alleles of almost each gene will exist, but if sequence similarity between alleles is high, the assembly process will usually lead to collapsing of allelic regions and only one allele will be present in the predicted gene and - in particular - protein set. This can be different in evolutionary old parthenogens, where independent accumulation of mutation in the alleles lead to sequence divergence (Welch and Meselson 2000) or allopolyploidy after hybridisation where homologs are divergent enough to be independently assembled (see (Glover et al. 2016); (Krasileva et al. 2013)). We found an excess of proteins for each of the parthenogenetic *Panagrolaimus* species in over 1,600 clusters of orthologs (orthogroups). Using the gene-tree reconciliation approach implemented in GRAMPA (Gregg et al. 2017) on FastME (Lefort et al. 2015) trees built from these orthogroups, we tested if auto- or allopolyploidy was the source of an extra set of proteins in the parthenogens (Fig 1a). The most parsimonious placement for these proteins found by GRAMPA was as basal to a clade containing both parthenogenetic and sexual species (**fig. 1**). It is thus unlikely that the extra copies arose by whole genome duplication within the parthenogenetic lineage (autopolyploidy), where a position as sister group to each of the parthenogens with the sexual species as common outgroup would have been expected (**fig. 1**). The tree topology supports an allopolyploid origin for these extra gene copies in the parthenogens.

To assess ploidy levels in the parthenogens we examined read depth coverage of variants in *P.* sp. PS1159 and presence of homeologous loci in *P. davidi*. Read mapping of RNA-Seq data on the predicted coding sequences (CDS) and variant calling yielded the expected result for the amphimict *P.* sp. ES5 sequenced from a homozygous population (fig. 2; similar pattern in *P. superbus* not shown), and the hermaphrodite JU765, where heterozygous sites had a mode of variants present at 50%:50% (**fig. 2**). However, in the parthenogenetic taxa we observed a clear peak in minor allele frequencies at ~33%, consistent with the presence of three copies at these sites (**fig. 2**). Using a two sample Kolmogorov-Smirnov test we confirmed that the allele frequency spectra of the parthenogenetic and amphimictic *Panagrolaimus* species, as well as the parthenogenetic *Panagrolaimus* and androdioecious *Propanagrolaimus*, belong to two different distributions (see Supplementary Table 1). Mapping of genomic reads to the CDS, repeat-masked genomes, and un-masked genomes resulted in similar allele frequency profiles (data not shown). These data support triploidy in the parthenogenetic *Panagrolaimus* species.

**Figure 2.**
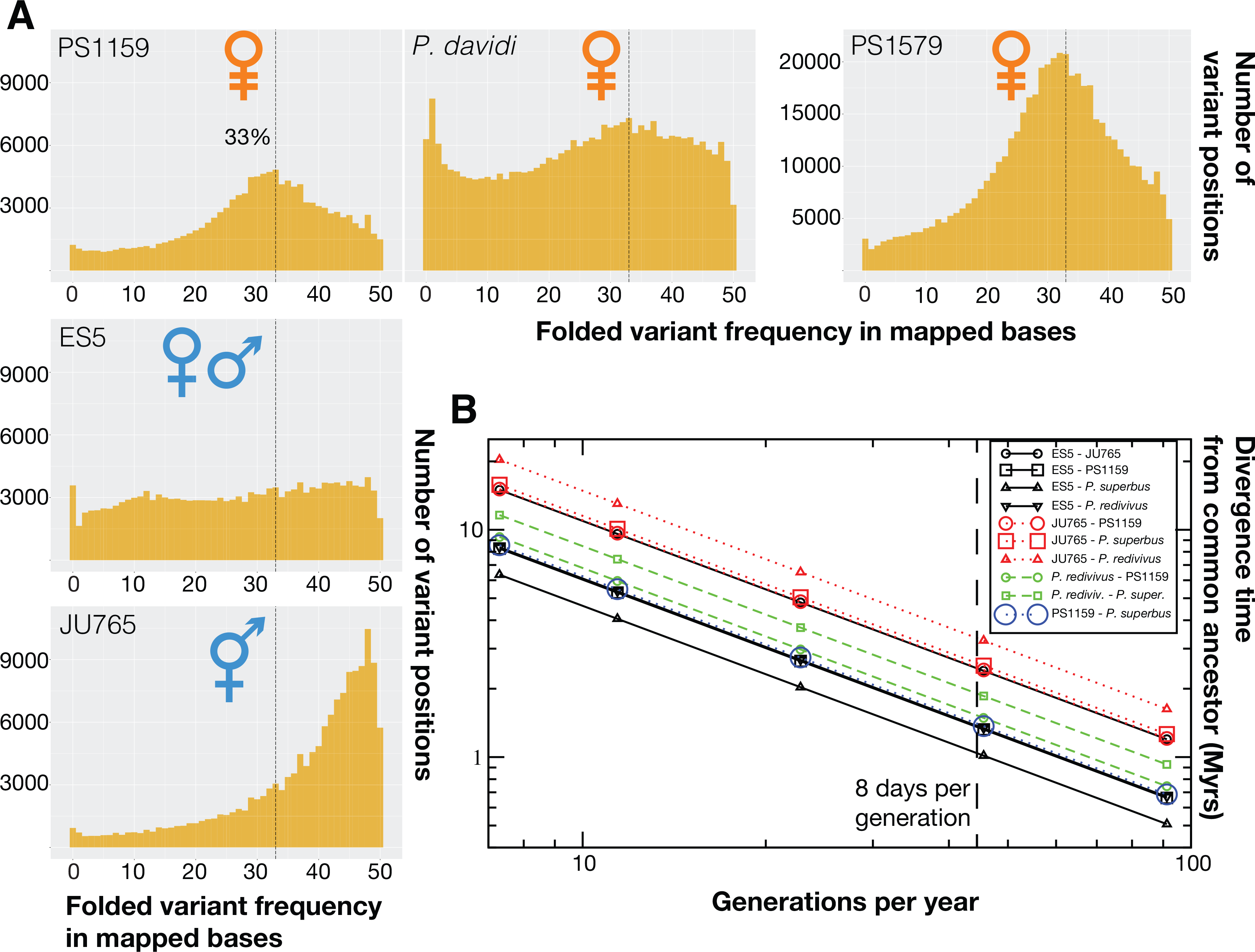
Polyploidy and hybridization analysis, and species age. **A.** Mapping RNA-Seq reads to coding sequences of predicted in the respective genomes and counting the occurrence of variant frequencies, we found that in the parthenogenetic species many variant positions are supported by 1/3 of the mapped reads at the position. Variant frequencies in the homozygous data from the gonochoristic ES5 are not showing a distinct pattern, while the hermaphrodite JU765 shows the 50% peak expected under heterozygosity. **B.** Using our own generation time measures, genome wide divergence calculated with Andi, and previous data for *Caenorhabditis* species as reference we calculated possible age of the origin of parthenogenesis in *Panagrolaimus*. 8 days per generation was measured in the laboratory and would indicate an approximate age of 1,3Myr for the PS1159/*P. superbus* split (see table 2).

Most parthenogenetic species are expected to be young. To estimate divergence time between the parthenogenetic and amphimictic *Panagrolaimus*, we calculated pairwise divergence of entire genome sequences with ANDI (Haubold et al. 2014). As calibration we used available divergence time estimates for *Caenorhabditis* species (Cutter 2008), corrected by generation time measurements in *Panagrolaimus* and *Propanagrolaimus*. The two amphimictic species *P. superbus* and *P.* sp. ES5 were the closest sisters (Table 1, **fig. 1**, also compare to supplementary fig 3). Assuming an average of 8 days per generation, the split between parthenogenetic and amphimicitc *Panagrolaimus* species is estimated to have occurred 1.3 - 1.4 Myr ago. Using a very conservative measure of 50 days per generation this split would have occurred 8 to 8.5 Myr ago.

**Table 1.** Pairwise divergence time estimates in *Panagrolaimus* spp. based on 8 days per generation (measured by us) and 50 days per generation as lower bound (see Supplement). Calibrated with divergence times calculated for *Caenorhabditis* spp. by (Cutter 2008). Mode of **repro**duction: **A**mphimictic, **H**ermaphroditic, **P**arthenogenetic.

| Species A | repro | Species B | repro | Myr (8 days/gen) | Myr (50 days/gen) |
| --- | --- | --- | --- | --- | --- |
| *P. superbus* | A | ES5 | A | 1 | 3.2 |
| PS1159 | P | ES5 | A | 1.33 | 8.25 |
| PS1159 | P | *P. superbus* | A | 1.37 | 8.55 |
| ES5 | A | *P. redivivus* | A | 1.35 | 8.35 |
| PS1159 | P | *P. redivivus* | A | 1.45 | 9.3 |
| *P. superbus* | A | *P. redivivus* | A | 1.88 | 11.6 |
| ES5 | A | JU765 | H | 2.4 | 14.95 |
| PS1159 | P | JU765 | H | 2.4 | 15.1 |
| *P. superbus* | A | JU765 | H | 2.5 | 15.8 |

### Genomic consequence of triploidy and parthenogenesis

The evolution of parthenogenesis through hybridisation is rapid. Newly formed hybrids may undergo genome and transcriptome shock, as co-adapted complexes can be disrupted by the new ploidy, and interacting partners that have accumulated differences through drift are brought into association. Such maladapted interactions need to be eliminated or moderated, and, in the case of parthenogens especially, the machineries of sex determination, fertilisation/oocyte activation and development are likely to undergo accelerated evolution. In parthenogens, the cost of producing males, which play no genetic part in species fitness is particularly high.

No males have been observed by us in any parthenogenetic *Panagrolaimus* species, not even under stress conditions (e.g. prolonged culturing at elevated temperatures or after reviving cultures from cryptobiosis). We therefore examined several genetic regulatory networks (GRNs) implicated in reproduction and sex in the parthenogens to identify changes consequent on the evolution of polyploidy and loss of sex. We carried out a strict orthology analysis using OrthoMCL (Li et al. 2003) with validation by Orthoinspector (Linard et al. 2014). Using GRNs defined by experimental intervention in *C. elegans* we identified homologous systems in the panagrolaims, and compared parthenogenetic and sexual *Panagrolaimus* species. Surprisingly, we found that many *C. elegans* genes acting in important developmental processes were absent not only from all *Panagrolaimus*, but also from other Tylenchina species. For example, in the sex determination GRN, tylenchine nematodes lacked orthologues of the *C. elegans* master regulator *xol-1*, as well as *fem-1, fem-3, her-1*, and other genes. Similarly, in the endoderm- and mesoderm-forming GRN tylenchines lacked *med-1, med-2, end-1, end-3*, and *spn-4*, amongst others. A similar pattern was observed for genes acting in spermatogenesis, early axis definition, DNA repair, and oogenesis. This core divergence from the systems as defined in *C. elegans* makes it difficult to identify changes associated with divergence in reproductive mode and ploidy.

While the parthenogenetic *Panagrolaimus* species had an excess of genes, many duplicated, in comparison to the amphimictic ones (**fig. 1**), the majority of genes were not present as duplicates. This is to be expected for two reasons: firstly, in the majority of cases the allelic copies are too similar and have been collapsed during the genome assembly process, and secondly, the most common fate of duplicated loci is loss through mutagenic inactivation. While the first, technical, reason is evidenced by the low genome assembly sizes (in comparison to the measured sizes) of the parthenogenetic panagrolaims, we wanted to test the second assumption. To this end, we identified clusters of orthologues that contained exactly one protein in *C. elegans*, in the two amphimictic, and all 3 parthenogenetic *Panagrolaimus* species (1:1:…:1 orthologues), as well as clusters where the parthenogenetic *P.* sp. PS1159 had more proteins than the amphimictic *P.* sp. ES5. Using the database of *C. elegans* RNAi phenotypes on wormbase.org we could retrieve phenotypes for 172 of the 703 single-copy one-to-one orthologues and 695 phenotypes for the 1,618 clusters with a higher number of proteins from the parthenogenetic species. A chi-square test confirmed (p-value = 0.0015) that the proportion of 91 lethal phenotypes (53%) in the 1:1 orthologues was significantly different from the proportion of 275 (40%) lethal phenotypes in the second group of clusters. This result is in accordance with the expectation that essential genes are less tolerant to copy number variation and changes in dosage.

### Horizontal gene transfer into panagrolaim genomes

We identified potential horizontal gene transfers (HGTs) using Alienness (Rancurel et al. 2017) (AI) approach. We identified from 22 (*P. redivivus*) to 232 *(P.* sp. ES5) likely HGT candidates from non-metazoan donors into the genomes of panagrolaimids (Table 2, Supplementary Excel files HGT AI). We used a series of additional screens to confirm HGT and rule out contamination, including a check for spliceosomal introns, codon usage, presence of evolutionarily conserved metazoan genes on the same contigs, and transcriptional support from RNA-Seq data. The inferred source taxa for the majority of high confidence HGT candidates were Bacteria followed by Fungi in all *Panagrolaimus* species except *P. davidi* (in which Fungi were the most frequent, followed by Bacteria).

We analysed OrthoMCL clusters composed of candidate HGT proteins that were either Panagrolaimidae-specific or shared with only one close outgroup. Twenty-six of these groups contained only one *Panagrolaimus* species and thus could represent either species-specific HGT events, or more ancient HGT that were secondarily lost. The remaining 110 OrthoMCL groups contained at least two *Panagrolaimus* species. We reconstructed the relative timing of acquisition *via* HGT for the 110 orthogroups using parsimony (Methods, **fig. 3**). Most (102; 93%) HGT events are predicted to have taken place in an ancestor of two or more *Panagrolaimus* species. The highest number of acquisitions (49 events) took place in the lineage leading to the last common ancestor of all the cryptobiotic *Panagrolaimus* species. Only seven HGT orthogroups were present in the last common ancestor of all eight panagrolaimid species analysed. These 7 ancestral HGT orthogroups plus the 49 novel acquisitions, suggests 56 independent HGT were present in the common ancestor of the cryptobiotic Panagrolaimus species. Eight orthogroups appeared to derive from multiple independent acquisitions at leaves of the tree. One HGT was predicted to have been acquired twice independently, at two nodes, but this may be a case of frequent loss.

**Table 2.**
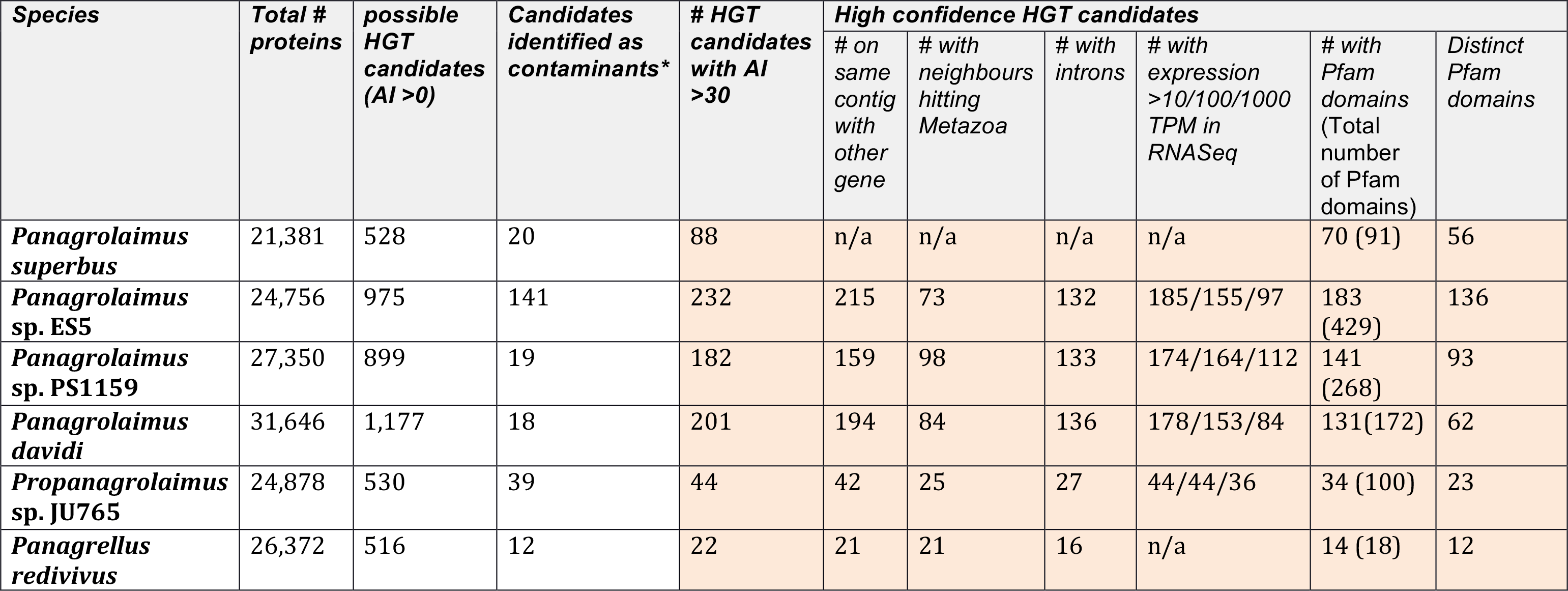
Protein numbers, HGT candidates, and high confidence HGT candidates in the *Panagrolaimus* genomes and outgroups. *AI>0 and >70% id to non-metazoan protein

| Species | Total # proteins | possible HGT candidates (AI >0) | Candidates identified as contaminants* | # HGT candidates with AI >30 | High confidence HGT candidates |  |  |  |  |  |
| --- | --- | --- | --- | --- | --- | --- | --- | --- | --- | --- |
|  |  |  |  |  | # on same contig with other gene | # with neighbours hitting Metazoa | # with introns | # with expression >10/100/1000 TPM in RNASeq | # with Pfam domains (Total number of Pfam domains) | Distinct Pfam domains |
| *Panagrolaimus superbus* | 21,381 | 528 | 20 | 88 | n/a | n/a | n/a | n/a | 70 (91) | 56 |
| *Panagrolaimus* sp. ES5 | 24,756 | 975 | 141 | 232 | 215 | 73 | 132 | 185/155/97 | 183 (429) | 136 |
| *Panagrolaimus* sp. PS1159 | 27,350 | 899 | 19 | 182 | 159 | 98 | 133 | 174/164/112 | 141 (268) | 93 |
| *Panagrolaimus davidi* | 31,646 | 1,177 | 18 | 201 | 194 | 84 | 136 | 178/153/84 | 131(172) | 62 |
| *Propanagrolaimus* sp. JU765 | 24,878 | 530 | 39 | 44 | 42 | 25 | 27 | 44/44/36 | 34 (100) | 23 |
| *Panagrellus redivivus* | 26,372 | 516 | 12 | 22 | 21 | 21 | 16 | n/a | 14 (18) | 12 |

The 910 proteins in 56 orthogroups putatively acquired via HGT in the six *Panagrolaimus* species were annotated with 192 different Pfam domains (Table 2). Only three Pfam domains were identified as conserved in the candidate HGT sets of all six *Panagrolaimus* species and all corresponded to enzymatic functions (alcohol dehydrogenase GroES-like domain, pyridine nucleotide-disulphide oxidoreductase and zinc-binding dehydrogenase) (Supplementary Excel file HGT - PFAM). HGTs conserved between *P. superbus, P.* sp. ES5, *P. davidi* and *P.* sp. PS1159 were annotated with thirty conserved Pfam domains that included GH43, GH28 and GH32 glycoside hydrolases, which have been identified as loci acquired by HGT in other nematodes (Haegeman et al. 2011; Rancurel et al. 2017).

### Loci potentially implicated in cryptobiosis in *Panagrolaimus*

We found that 47 Pfam domains functionally connected to cryptobiosis were associated to proteins putatively acquired via HGT. Some of these were acquired at ancestral nodes: 19 at the last common ancestor of *Panagrolaimus* and one predicted to have been present in the tylenchine ancestor. Five were specific to all or several parthenogenetic species. Functions associated with these domains included peptidase, diapausin, killer toxins of the Kp4 family, LURP-one-related proteins, flavin-binding monooxygenase-like protein, caleosin, and photolyase (see Supplement for extended description). Proteins with these functions fulfil various roles in defence against pathogens and the removal of xenobiotics (Gu et al. 1995; Kuwabara et al. 1999; Tanaka et al. 2003; Eswaramoorthy et al. 2006; Khalil et al. 2014). The photolyase (**fig. 3**) was of particular interest, since photolyases are able to repair UV induced damage in DNA (Sancar 1990) and have been lost in several branches of Metazoa (Lucas-Lledo and Lynch 2009), and genome repair is important for desiccating organisms (Jönsson and Schill 2007). The caleosin found in *Panagrolaimus* is the first report to date of animal caleosins, since these proteins have previously been described only in multicellular plants, green algae, and fungi (Partridge and Murphy 2009). Plant caleosins expressed in non-seed tissues are responsive to a range of environmental stresses, particularly salinity and dehydration (Blée et al. 2014).

**Figure 3.**
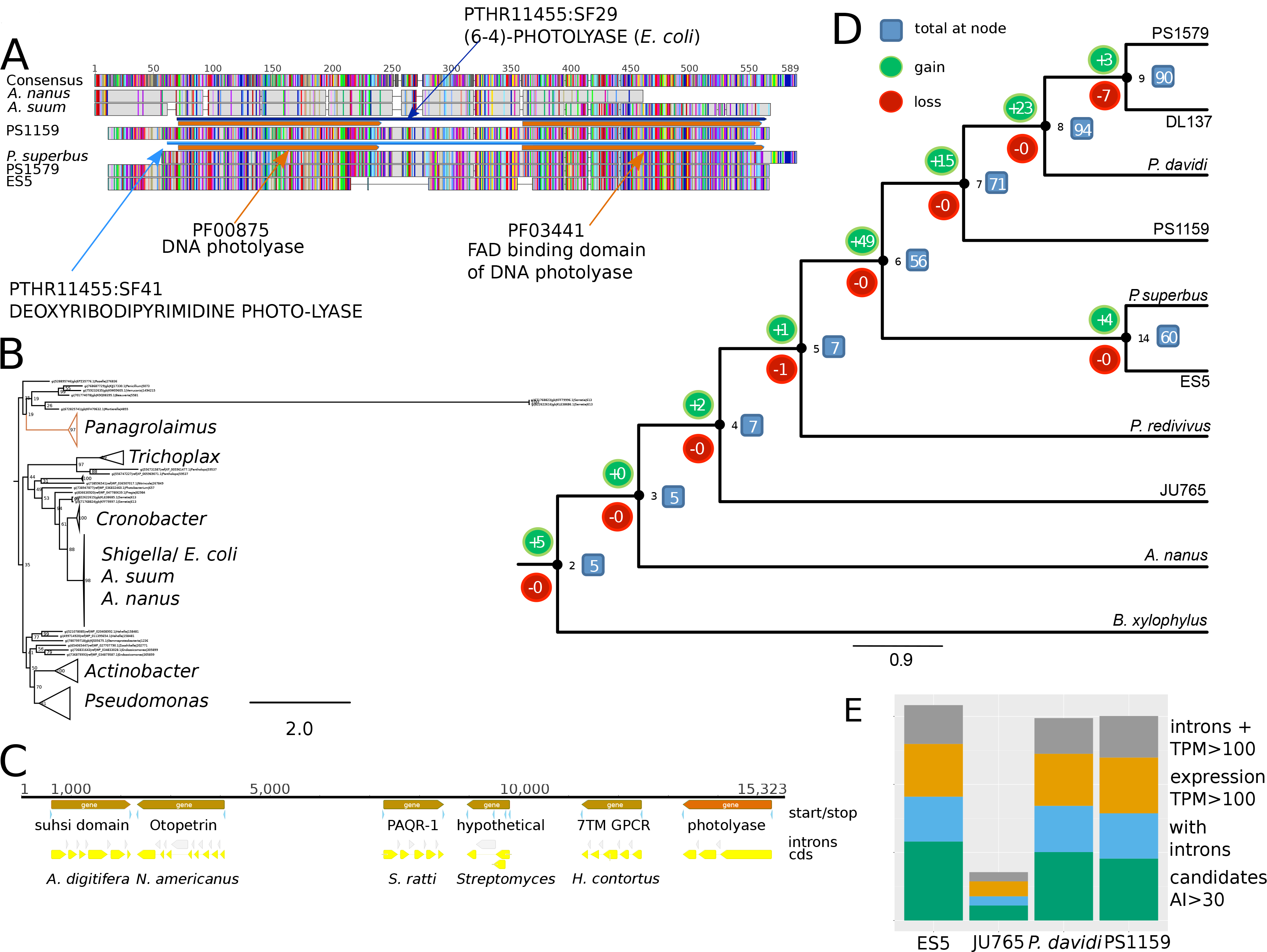
Reconstruction of the timing of acquisition and loss of HGT candidates. **A.** An alignment of the photolyase found in *Panagrolaimus*, and *A. suum* and *A. nanus* shows that distinct differences between the proteins. The *A. suum* and *A. nanus* domain annotations point towards *E. coli*, rendering it likely that the proteins are contaminations from the bacterial food source used commonly used in nematode cultures. **B.** A phylogenetic tree of the photolyase proteins in Panagrolaimus, A. *suum* and *A. nanus,* and bacterial photolyases identified with blast confirms that the *A. suum* and *A. nanus* proteins are *E. coli* contamination, while the *Panagrolaimus* proteins have evolved since their horizontal acquisition. **C.** Demonstrating the integration into the *Panagrolaimus* genomes the photolyase was found on one contig with eukaryotic (nematode) proteins in the *P. sp.* PS1159 assembly. **D.** Cladogram of the Panagrolaimidae species and two outgroup relatives. The distribution of HGT candidates across these species has been mapped to the phylogeny and ancestral states have been deduced with a parsimony approach. Internal nodes are numbered in black (2,3,4,5,6,7,8,9,14). The number of gene families acquired via HGT / lost at each node are represented in green / red bubbles, respectively. The total number of gene families acquired via HGT present at each node is indicated within blue boxes. **E.** Stacked bars of classes within the AI>30 HGT candidates for the different species. Candidates with introns and expression levels of TPM>100 can be regarded as most reliable.

Adaptation to novel habitats can also involve expansion and diversification of gene families. We therefore analysed gene family expansions in the cryptobiotic *Panagrolaimus* species compared to related taxa. Forty-three Pfam domains were overrepresented in *Panagrolaimus* compared to an outgroup containing Prop. sp. JU765 and *anagrellus redivivus*, after Benjamini-Hochberg correction, and 123 Pfam domains were overrepresented in Panagrolaimidae (*Panagrolaimus, Propanagrolaimus*, and *Panagrellus*) compared to the remaining taxa (**fig. 4**).

We employed a support vector machine (SVM) classifier based on Pfam domain annotations to distinguish *Panagrolaimus* from *Propanagrolaimus* plus *Panagrellus*, and, independenly Panagrolaimidae from the outgroup. Classification performance was significantly better than control classification based on a permuted class-sample mapping in the training set (real *versus* permuted classifier performance in the *Panagrolaimus versus Propanagrolaimus/Panagrellus* SVM: *Panagrolaimus:* T = 7, *Propanagrolaimus/Panagrellus:* T= 7.4, real *versus* permuted classifier performance in the Panagrolaimidae *versus* outgroup SVM: Panagrolaimidae: T = 36.9, outgroup: T= 39, all p < 0.001, all df = 1999). In the *Panagrolaimus versus Panagrellus/Propanagrolaimus* SVM analyses, 64 domains were frequently associated with successful classification, of which 13 were common to the SVM and the two-sided Fisher’s exact test approaches (**fig. 4**). These 13 domain annotations included several that may play a role in cryptobiosis, including Hsp70 protein domains, serine protease inhibitors, ubiquitin family domains, lipase domains and enoyl-reductase domains. Annotations found to be overrepresented only in the two-sided Fisher’s exact test analyses were proteolysis (subtilase family and matrixin domains) and maintenance of genome integrity (PIF1-like helicase, deoxyribonuclease II, linker histone H1 and H5 family).

In the SVM comparison of Panagrolaimidae *versus* outgroup species 80 domains were found to contribute to the differentiation between the taxa. Of these 24 were common with those identified in the two-sided Fisher’s exact test analyses (**fig. 4**), including BTB/POZ domain, BTB and C-terminal Kelch, and ABC transporter. These three domains had previously been shown to be expanded in *P. redivivus* compared to *C. elegans (Srinivasan et al. 2013)*. While C-type lectins were overrepresented in Panagrolaimidae as a whole, in the two-species comparison *C. elegans* had more C-type lectins than did *P. redivivus (Srinivasan et al. 2013).* C-type lectins are expressed during dehydration in the tylenchine nematode *Aphelenchus avenae (Reardon et al. 2010)* and could thus also play a role in the dehydration process in *Panagrolaimus*. Other domains with potential roles in cryptobiosis were also identified as amplified in Panagrolaimidae, including heat shock proteins (HSPs), such as HSP70 and the collagen-specific serpin HSP47, the late embryogenesis abundant (LEA) domain (identified only in the NHST analysis), and antioxidant and reactive oxygen species (ROS) regulating domains such as cytochrome P450. The abundance of the DEAD/DEAH box domain, involved in RNA metabolism, may support roles in transcript and ribosome stability (Guan et al. 2013). Small heat shock proteins (sHSPs) carry out chaperone functions in anhydriobiotic animals (Gusev et al. 2011). We identified a tylenchine-specific family of sHSPs that was distinct from the lineage specific expansion in *C. elegans (Aevermann and Waters 2008)*, but no subset specific to panagrolaims (Supplementary Figure 2).

**Figure 4.**
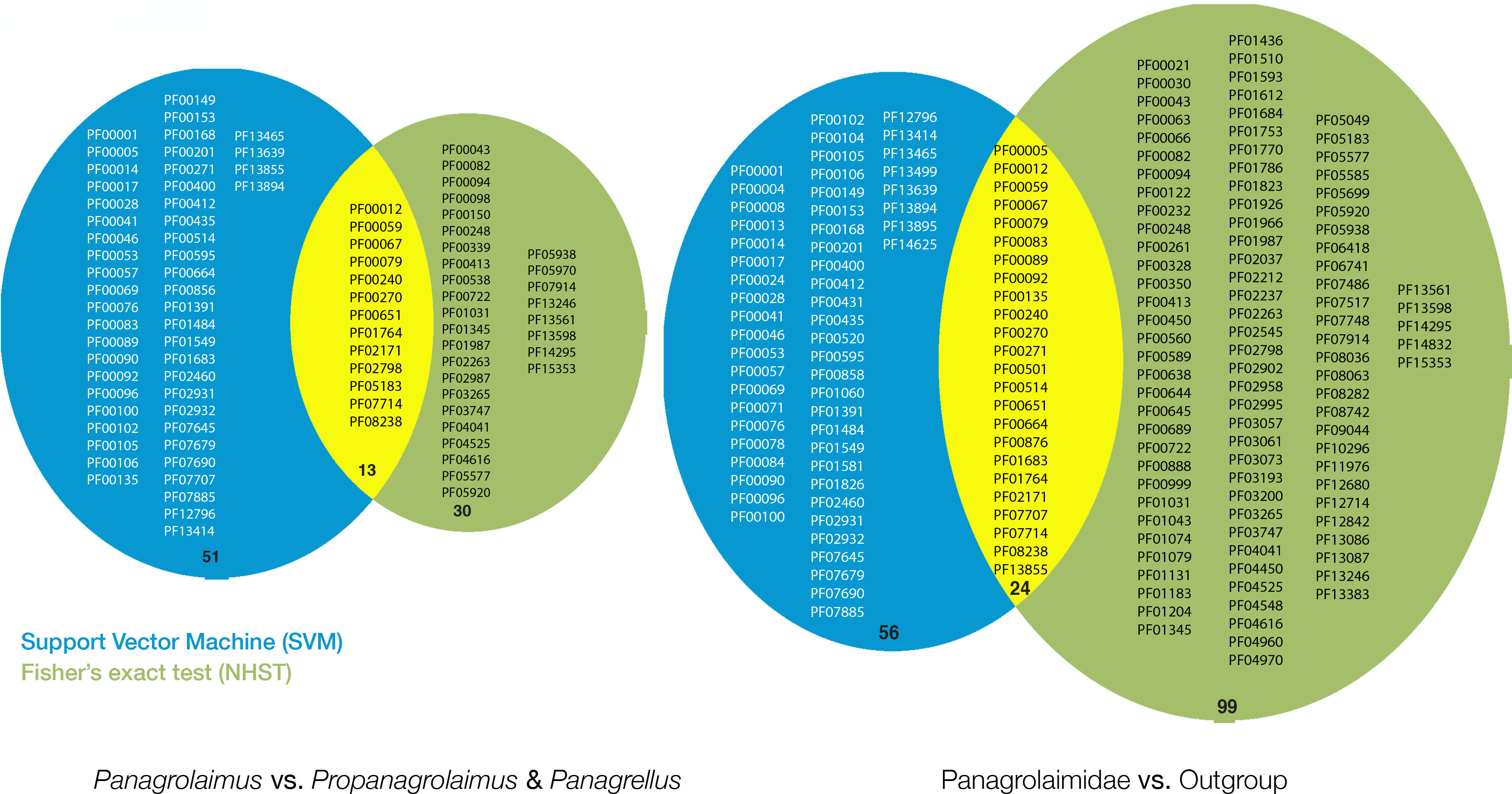
SVM classification and overrepresentation analysis. Using a Fisher’s exact test and a Support Vector Machine classification we compared Pfam annotations in the genus *Pangrolaimus* with thos of *Propanagrolaimus* and *Panagrellus* (combined), as well as these species taken together as Panagrolaimidae with outgroup species. This comparison revealed several protein families associated with cryptobyosis, e.g HSPs, helicases, and C-type lectins, to be important for *Panagrolaimus* biology.

### Loci under positive selection in *Panagrolaimus*

Measuring dN/dS ratios we identified candidate genes under positive selection comparing species with different reproductive modes and comparing cryptobiotic with non-cryptobiotic panagrolaims (**fig. 5**). InterProScan and GO annotation of these loci showed significant enrichment of functions such as carbohydrate transport and metabolism, cell wall/membrane/envelope biogenesis, and replication, recombination and repair (**fig. 5**, Supplementary Excel file dNdS). Loci contributing to “replication, recombination and repair” annotations include several connected to DNA integrity and processing of nucleic acids (**fig. 5**). An orthologue of *C. elegans denn-4*, which is expressed in neurons and also involved in reproduction, was possibly under positive selection in parthenogenetic compared to amphimictic *Panagrolaimus*. Homologues of the human DNA polymerase zeta (REV3L) were under positive selection. These might act in DNA repair, maintaining genome integrity in the nematodes. *Panagrolaimus* orthologues of *C. elegans fcd-2*, which acts on cross-linked DNA and DNA damage repair (Vermeulen 2015), also had a dN/dS >1 between *Panagrolaimus* and the outgroups, as did helicases similar to *C. elegans* WBGene00010061. In comparison to the parthenogenetic species *fcd-2* also has a signature of selection in the sexual species. It may therefore play a role in cryptobiosis, as the parthenogenetic species are less amenable to desiccation than the sexual congeners (McGill et al. 2015). Similarly, the *Panagrolaimus* orthologue of the *C. elegans* fatty acid and retinol-binding protein *far-6* (Garofalo et al. 2003) appeared to be under positive selection. Fatty acids and retinol are required for the biosynthesis of lipids integral to the nematode cell membranes and cuticle. Thus, effective lipid binding and transport may have a significant role in the evolution of cryptobiosis.

**Figure 5.**
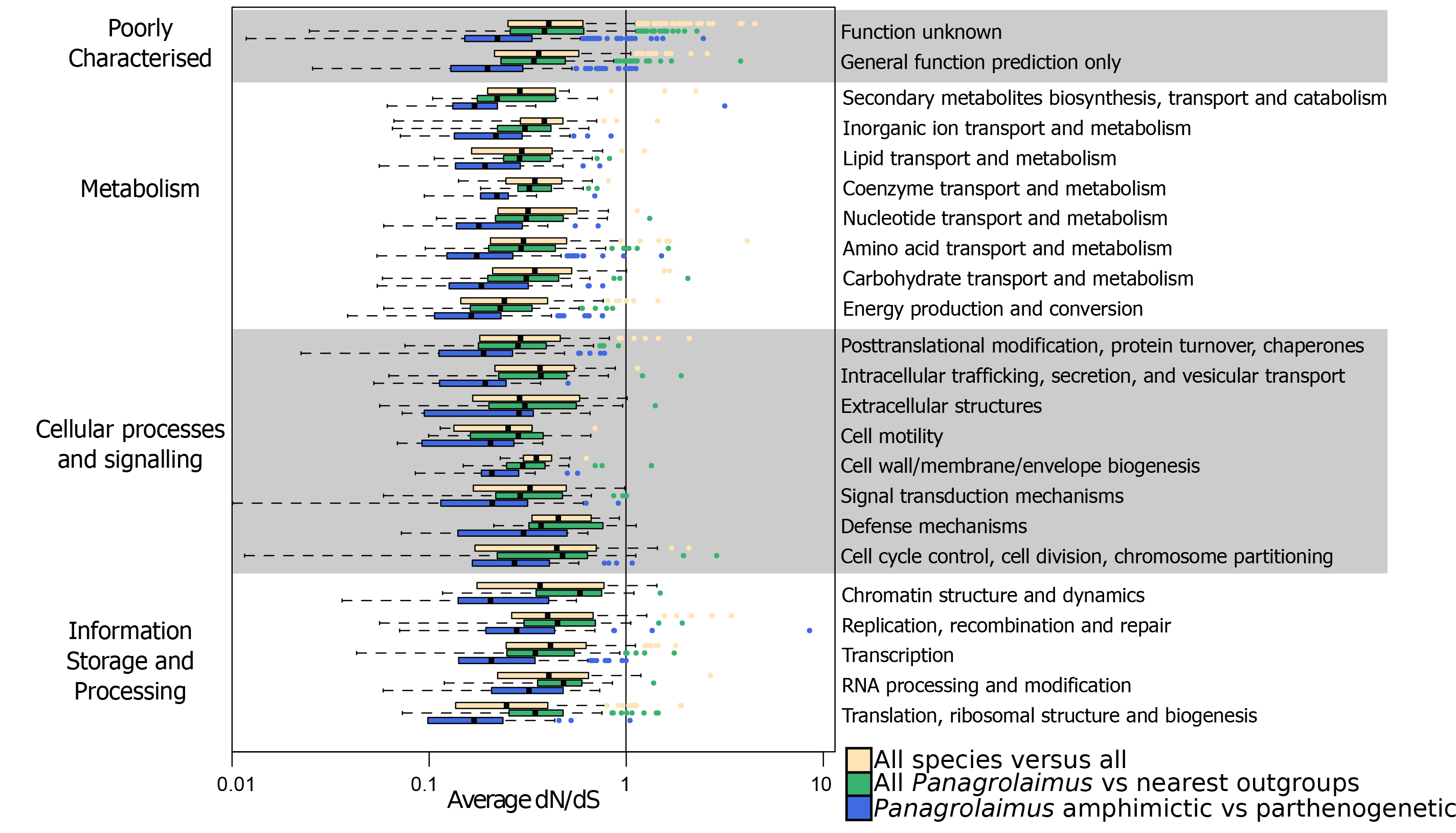
Adaptive evolution analysis. Average rates of evolution (dN/dS) are shown for comparisons between parthenogenic and gonochoristic *Panagrolaimus* (in blue), *Panagrolaimus* and the closest outgroups (*Propanagrolaimus* and *Panagrellus*) (in green), and between all species (in yellow) grouped by functional categories (see Supplementary Excel file dNdS). We find that while the majority of genes have a ratio of < 1, there are notable exceptions across several functional categories both between parthenogenetic and gonochoristic species and between panagrolaimids and their closest relatives.

## Discussion

Understanding the evolution and persistence of parthenogenesis is fundamental for answering why sex is such an ubiquitous mode of reproduction in animals. A similarly important question is how animals adapt to extreme environmental conditions. It has been proposed that both traits are linked in geographical parthenogenesis.

Many parthenogenetic nematode species are of hybrid origin and polyploid (Castagnone-Sereno and Danchin 2014; Lunt et al. 2014; Hiraki et al. 2017; Kraus et al. 2017); (Lunt 2008; Lunt et al. 2014; Blanc-Mathieu et al. 2017). It is a classical tenet of evolutionary biology, that polyploid animals, in particular heterogametic organisms, have a reduced fitness or are not viable (Haldane 1922; Burke and Arnold 2001). In contrast, Lynch argued that polyploidy might under certain circumstances lead to a fitness advantage in buffering against Muller’s ratchet (Lynch 1984). Hybridisation leads to aberrant meiosis in animals, and a possible escape from this dilemma could be parthenogenesis (Avise 2008). In such systems, incorporation of a round of chromosomal endoreplication is a possible route to regain triploidy during oogenesis (Lutes et al. 2010).

We have analysed the genomes and transcriptomes of parthenogenetic and amphimictic *Panagrolaimus* nematodes, which in contrast to the outgroup *Propanagrolaimus* and *Panagrellus* are both capable of surviving under prolonged anhydro- and cryobiosis. Our combined karyotypic and molecular data indicate a triploid hybrid origin in parthenogenetic *Panagrolaimus*, with all analysed parthenogenetic species showing n=12 (compared to diploid n=8 in amphimicts), and having an excess of duplicate genes that share a single phylogenetic origin with variant frequency spectra compatible with triploidy. Triploidy could have arisen by autopolyploidy (duplication of one set of ancestral chromosomes, for example by fertilisation of a diploid oocyte by a haploid sperm from the same species) or allopolyploidy (where the additional haploid complement comes from a distinct lineage). These contrasting origins in turn predict contrasting patterns of divergence between the three copies of each locus in a triploid: the autopolyploid will have three closely related loci, while the allopolyploid will have a pair of closely related (allelic) copies and a more distantly related (homoeologs) copy. In meiotic *Panagrolaimus* parthenogens we found - similar to what was observed in mitotic *Meloidogyne* parthenogens (Blanc-Matthieu et al. 2017) - phylogenetic evidence for divergence of extra copies, and probable acquisition from a donor species positioned between the *Propanagrolaimus* and *Panagrolaimus* species present in our analysis. These results support allotriploidisation.

Analysis of different taxa that can undergo anhydrobiosis has identified several different kinds of genes associated with desiccation survival. In plant embryos, LEA proteins, natively unstructured peptides that are believed to surround proteins like a “molecular shield” (Chakrabortee et al. 2012), are highly expressed. Natively unstructured proteins have been identified in anhydrobiotic responses in nematodes (*Aphelenchus avenae (Reardon et al. 2010)*), arthropods (*Polypedilum vanderplanki (Cornette et al. 2010)*) and tardigrades (*R. varieornatus* and *H. dujardini (Yoshida et al. 2017)*). Other gene families associated with anhydrobiosis include protein kinases, proteasomal components, ubiqutin, protease inhibitors, proteases, DNA repair enzymes, enzymes with roles in oxidative stress protection, and heat shock proteins (Cornette et al. 2010). We identified overrepresentation of the above-mentioned gene families, as well as a number of others not previously identified, associated with cryptobiotic *Panagrolaimus* species. The set of *Panagrolaimus*-enhanced activities overlapped with the list of genes we identified as having been acquired through HGT. Many putative HGT were mapped to the root of the cryptobiotic *Panagrolaimus* species group. These candidate HGT genes have acquired spliceosomal introns, have similar GC content and codon usage to resident *Panagrolaimus* genes, are assembled in the vicinity of *bona fide* nematode genes and are expressed as poly(A)+ RNA. They can be regarded as firmly domesticated and functionally integrated into the host genomes. Functional annotation of these putative HGT loci suggested that they contribute to anhydrobiotic physiology. Genes acquired through HGT were also implicated in cryptobiosis in the rotifer *A. vaga* and the tardigrade *R. varieornatus* (Yoshida et al. 2017), and a functional link between anhydrobiosis and HGT has been proposed (Hespeels et al. 2014).

The theory underpinning geographical parthenogenesis holds that parthenogenetic species have an advantage in adapting to new and/or extreme environments. Since we did not observe a signal for specific adaptations in the parthenogenetic species (including to the mode of reproduction itself) in comparison to sexual congeners it appears possible that a general-purpose genotype is indeed maintained in evolutionary young parthenogens, potentially by combining several differently-adapted genomes during hybridisation. Estimates of generation times cannot be easily transferred from the laboratory to nature, and for example the Antarctic species *P. davidi* could potentially be restricted to very few generations per year. However, other species are present in temperate regions and thrive in semi-arid (sandy) soils or leaf litter (Shannon et al. 2005) (McGill et al. 2015) where many generations per year are likely. Based on this, our estimate that the *Panagrolaimus* species, like most parthenogens, are younger than the ancient evolutionary “scandals” (Smith 1978) are credible.

Few parthenogenetic species are suitable for genomic and molecular research. *Panagrolaimus* nematodes are easily cultured in the laboratory and accessible to molecular genetic analysis (Schiffer et al. 2014), including gene knock-out techniques (Reardon et al. 2010). *Panagrolaimus* species may also be useful in understanding developmental diversity in Nematoda, diversification and speciation in parthenogens, the effects of triploidy on gene and genome evolution, and the biology of cryptobiosis. The existence of a well-studied model species (*C. elegans*) as a comparator should make the *Panagrolaimus* model even more attractive, even though the evolutionary distance between these species precludes some direct transfers of information. Our exploration of the genomes of different Panagrolaimid species has revealed the first possible origins and evolution of a cryptobiotic biology in nematodes. The genome analysis also highlighted and reinforced some singularities as signatures of an allopolyploid and parthenogenetic lifestyle.

## Materials and Methods

### Nematode strains and culture

Nematodes were acquired from collaborators or through sampling by one of us (E.S.) and cultured in the laboratory on low-salt agar plates (Lahl et al. 2003). Other tylenchine nematode genomes analysed were obtained from WormBase (www.wormbase.org).

### Genome sizes and chromosome numbers

We estimated genome sizes in several isolates of *Panagrolaimus* using Feulgen image analysis densitometry (FIAD, following the protocols in (Hardie et al. 2002)), integrated optical densities (IODs) from at least 14 nuclei per individual worm were compared with erythrocytes of *G. domesticus* (C-value =1.25 pg) under x100 magnification (immersion oil, nD = 1.5150). The DNA content in picograms was converted to megabases using the formula 1 Mb = 1.022 × 103 pg or 1 pg = 978 Mb (Dolezel et al. 2003). These measurements were complemented by RT-D-PCR assay (Wilhelm et al. 2003) based on orthologues of *C. elegans* locus ZK682.5 (WBGene00022789), which has been identified as a nematode-wide single copy gene (Mitreva et al. 2011). Chromosomes were counted, after DAPI staining, in oocytes and 1-cell embryos. For this, gravid nematodes were picked from plates and transferred to M9 buffer and dissected with an insect needle to release gonadal tubes and embryos. These were transferred with a capillary tube into a small drop of 2μg/ml DAPI (Sigma) on a microscope slide, squashed under a coverslip in the stain and the preparation was sealed with nail varnish. After 10-15 min staining the preparations were analysed under an Olympus FluoView1000 confocal microscope with a 60x (NA 1.35) oil objective. Karyotypes were taken as valid only when confirmed in 10 independent chromosome sets. See supplementary fig. 1.

### Genome Assembly

We constructed genome assemblies from Illumina short read libraries. All libraries are deposited in the SRA under the BioProject PRJNA374706. After running first pass assemblies with the CLC Assembly Cell v.4.2 we applied the khmer pipeline (Brown et al.) to digitally normalise read coverage on ES5, *P. superbus*, JU765 and PS1159 data. We identified and removed contaminating bacterial sequences using the blobtools approach (Kumar et al. 2013). The cleaned reads were reassembled using the Velvet assembler (Zerbino and Birney 2008) exploring different k-mer sizes. We then employed RNA-Seq derived mRNA predictions (see below) to scaffold the genomes with the SCUBAT pipeline (https://github.com/elswob/SCUBAT). For *P. superbus* the genome input to SCUBAT originated from a CLC assembly, as this proved better than the tested Velvet assemblies. For *P. davidi* DAW1, where the original assembly (Thorne et al. 2014) was of low completeness (39% complete KOGs as assessed by CEGMA^8^), we (re)assembled the genomes with SPAdes (Bankevich et al. 2012) and the redundans pipeline (Pryszcz and Gabaldón 2016). We also used SPAdes and the redundans pipeline to assemble the *P.* sp. PS1579 genome. Assembly qualities were evaluated with custom scripts. See supplementary Fig 3 for CEGMA completeness. We utilized GNU Parallel (Tange 2011) in various steps of the assembly pipeline and in downstream analyses.

### Gene Prediction

We used Augustus (Stanke and Waack 2003) (v.3.0.1) for gene prediction, employing an iterative approach to generate the most credible predictions. Augustus was trained with CEGMA-predicted genes. Wherever RNA-Seq data were available (*P.* sp. ES5, *P.* sp. PS1159, *Prop.* sp. JU765), it was assembled based on alignment to the genome, using gsnap (Wu and Nacu 2010), which has been shown to perform well in finding correct splice sites (Sipos et al. 2013). These mappings then served as evidence for Augustus in a second round of gene prediction. For the first round of Augustus training in *P. superbus* we used exonerate (Slater and Birney 2005) to align predicted *P. sp.* PS1159 proteins, as no RNA-Seq data were available. We then mapped findorf (Krasileva et al. 2013) predicted ORFs (see below) from the *P. superbus* transcriptome using BLAT (Kent 2002) and created Augustus evidence from this. For *P. davidi* no transcriptomic data were available at the time of our analyses, and we used the *P.* sp. PS1159 species model in Augustus along with the *P. davidi* CEGMA genes as hints. We implemented a best hit BLAST approach using different taxonomic databases to identify likely contamination in predicted genes.Candidate contamination was removed from the predicted gene sets and from the corresponding contigs in the genome assemblies. In the ES5 genome we identified an excess of *E. coli* contamination and employed mugsy (Angiuoli and Salzberg 2011) for full genome alignments between the nematode and *E. coli* to identify and remove the corresponding contigs. Contamination was also additionally screened for in the HGT assay (see below).

Open reading frames were predicted from *de novo* transcriptome assemblies (*P. superbus*, DL137, PS1579) using findorf. The findorf approach involved comparison of transcripts with proteomes from other nematode species (*Brugia malayi*, *Caenorhabditis briggsae, Caenorhabditis elegans, P.* sp. ES5, *Prop.* sp. JU765, *Meloidogyne hapla, Panagrellus redivius, P.* sp. PS1159, and preliminary data from *Plectus sambesii*) using BLASTX to identify frameshifts and premature stop codons, and Hidden Markov models (HMMs) to identify Pfam domains to infer ORF start positions. In total, findorf inferred 21,381 *P. superbus* ORFs, 34,868 *P.* sp. DL137 ORFs and 47,754 *P.* sp. PS1579 ORFs.

### Proteome Annotation

Putative domains in 13 nematode proteomes were inferred using InterProScan (Jones et al. 2014) version 5.7-48.0. We analysed all our *Panagrolaimus* species (including both transcriptomic and genomic data for *P. superbus*), our *Propanagrolaimus* outgroup, and the two remote outgroup species *Caenorhabditis elegans* and *Ascaris suum.* A summary of the Pfam domains annotation is deposited on the accompanying genome hubs webpage for the *Panagrolaimus* genomes.

### Orthologous Proteins

We inferred orthology between proteins from our newly sequenced panagrolaimids and a set of outgroups (other species from the Tylenchina - Clade IV of Nematoda *sensu* Blaxter (Blaxter et al. 1998)) using OrthoMCL (v.2.0.8) (Li et al. 2003), Orthoinspector (v.2.11) (Linard et al. 2014), and Orthofinder (Emms and Kelly 2015). We also included preliminary data from the cephalobid species *Acrobeloides nanus* (courtesy of Itai Yanai, NYU, New York), as well as the model *C. elegans* (Rhabditina; Clade V) and *Ascaris suum* (Spirurina; Clade III). To detect orthologs to key *C. elegans* developmental genes (see section on GRNs) we primarily relied on the extensively tested and robust OrthoMCL pipeline based on NCBI blast, which we validated with Orthoinspector. The latter found additional divergent orthologous, which were hidden to OrthoMCL. Orthofinder based on Diamond (Buchfink et al. 2014) BLAST was only used in the GRAMPA (Thomas et al. 2016) analysis, since the output contains gene trees for each cluster (see section on ploidy below). Were OrthoMCL and Orthoinspector grossly disagreed or the presence/absence pattern across species appeared inconsistent in the GRN analysis, we directly screened genomes for the presence of a gene using the HMM profile figmop pipeline (Curran et al. 2014).

### Assessment of ploidy

We first mapped reads against the assembled small subunit ribosomal sequences in each species and identified variants. This suggested the presence of a significant “minor allelic variant” in the parthenogenetic species. Both the hermaphroditic species *Prop.* sp. JU765, expected to be diploid, and the potentially polyploid parthenogenetic strain *P.* sp. PS1159, were inbred through 30 generations of single offspring propagation, this way expecting largely homogenised genomes. In contrast, *P. davidi* was expected to be more heterozygous as large populations from multiple plates were used for DNA extraction. We extracted all Augustus predicted genes for each species and mapped RNA-Seq read sets against the coding sequences using the CLC mapper (v.5.0), requiring a sequence identity of 90% and a read length threshold of 80%. We performed the same mapping approach with genomic reads against repeat masked and unmasked genomes. We used BamTools (Barnett et al. 2011) (v.2.3.0) to sort the mappings, SAMtools (Li et al. 2009) (v.1.0-13) to create mpileups, VarScan (Koboldt et al. 2012) (v.2.3.6) to call variants, and collated the minor variant frequency spectrum (folded site spectrum) for each species. We then implemented a two sample Kolmogorov Smirnov (KS) test available in the Julia (Bezanson et al. 2012) programming language to analyse whether the observed variant frequencies are different under the null hypotheses that the frequencies are sampled from the same distribution (Supplementary Table 1): we first sampled 20,000 data points from each distribution and directly compared these with the KS test; next we ran 50,000 bootstrap replicates on 20,000 randomly mixed samples per distribution, and then measured the exact p-value in the bootstrapped distribution.

For the GRAMPA (Thomas et al. 2016) analysis testing allo- vs. autopolyploidy we used all ~ 15k FastME (Lefort et al. 2015) trees from running Orthofinder mid-point rooted with NOTUNG (2.8.1.7) (Chen et al. 2000). To validate these results we also used a more comprehensive initial dataset including the DL137 and PS1579 proteomes and selected 2313 groups of orthologues from OrthoMCL that had a 2:1 ratio of parthenogenetic to sexual species. For each group we built an alignment with clustal-omega, inferred a protein tree with RAxML and ran GRAMPA. This resulted in the same topology as the Orthofinder FastME based GRAMPA tree shown in Fig 1.

### Estimation of divergence times

We used ANDI (Haubold et al. 2014) to calculate pairwise evolutionary distances between three species of *Panagrolaimus* (*P. superbus, P.* sp. ES5, and *P.* sp. PS1159) and two outgroups (*Propanagrolaimus* sp. JU765, and *Panagrellus redivivus*) (see Supplementary Table 2). We excluded the parthenogenetic *P. davidi* from this analysis since the assembly span of 118Mb and our variant calling (see above) indicate this assembly to contain duplicate regions when compared to the quasi-haploid assembly of PS1159 (85Mb). ANDI is designed for comparisons of entire genomes, and does not require homologous sequences or genes as input. To convert ANDI-distances into divergence measured in generations or in absolute time we used the divergence of *Caenorhabditis briggsae* and *Caenorhabditis* sp. 5 as a reference. The *Caenorhabditis* species are separated by about 171 M generations (i.e. 85.5 M generations since their last common ancestor (Cutter 2008)). Generation time measurements were conducted by Isabel Goebel at the University of Köln (unpublished “Staatsexamen” thesis “Untersuchungen zum Lebenszyklus von Nematoden mit unterschiedlichem Fortpflanzungsmodus”) under supervision of PHS and ES. Briefly, nematodes from parthenogenetic strains *P.* sp. PS1159, and *P.* sp. PS1579, and the sexual species *P. superbus, P.* sp. ES5, and *Prop.* sp. JU765 were cultured at room temperature on small *E. coli*-seeded agar plates. These were monitored every few hours during early development and then every day during adult life. Across all species assayed, generation times (Vancoppenolle et al. 1999) averaged 8 days under our laboratory conditions. Since there is some uncertainty in the generation length in nature we estimated timings based on 4, 8, 16, 32, and 50 days per generation (corresponding to around 91, 46, 231, 111 and 7 generations per year, respectively).

### Adaptive evolution

We used Crann (Creevey and McInerney 2003) to explore signatures of selection in panagrolaim genes. Over 20,000 OrthoMCL derived orthology cluster protein alignments, built using clustal-omega, were reverse-translated into DNA alignments with trimAl and SAMtools. We retained 7,335 alignments that contained at least one parthenogenetic and one amphimictic *Panagrolaimus* species. Further filtering of short and outlier sequences within each of these gene family alignments, using amino acid versions of the sequence alignments, removed poorly-defined clusters. If the length of any sequence was less than half the length of the overall alignment it was removed. For all remaining sequences in each alignment, two filtering steps were carried out: A distance matrix was constructed from the amino acid versions of the alignments using ProtDist from the PHYLIP (Felsenstein 1995) package and the JTT model of evolution was used to construct a neighbour-joining tree with neighbour from the PHYLIP package. Using custom scripts, the average taxon-taxon path length on the tree was calculated, and outlier sequences which had a significantly greater average path length were identified using the R statistical package. These sequences were removed from both the DNA and amino acid alignments. The second filter checked for saturation in the nucleotide versions of the alignment. This was carried out by building two trees with PAUP for each alignment, the first using a P distance and the second using the HKY model. The taxon-taxon length across both trees were calculated as before and both calculations combined for visualization as a scatter plot, using R. Any alignments exhibiting signatures of saturation following visual inspection of the plots were removed from both the DNA and amino acid alignments. The resulting filtered DNA alignments were input to Crann to calculate all pairwise rates of non-synonymous substitutions per non-synonymous site (dN) and synonymous substitution per synonymous site (dS). The average of all pairwaise dN/dS ratios between the following categories of species were calculated: (1) sexually-reproducing *Panagrolaimus* species versus parthogenetic *Panagrolaimus* species; (2) all *Panagrolaimus* species, versus nearest outgroups; (3) all species in analysis versus all. See supplementary fig. 5. The rates of dN/dS were sorted into categories by their functions (Tatusov et al. 1997) (fig. 4).

### Support vector machine classification

We parsed inferred InterProScan annotations with custom Python scripts, focusing on Pfam domains. Statistical over-representation (Fisher’s exact test) was estimated using custom Python scripts. We corrected for multiple testing using the Benjamin-Hochberg algorithm (for false discovery rate) implemented in the R (R Core Team) statistical language. We included transcriptomic data from *P.* sp. DL137 and *P.* sp. PS1159, as well as the *P. davidi* DAW1 gene predictions (which were expected to include duplications) in our test to retrieve proteins potentially missed in the genome-wide annotations of *P.* sp. PS1159 alone.

To improve on the classical null hypothesis significance testing (NHST), we employed computational classifier analysis. We employed a support vector machine (SVM) to address two pivotal questions: (1) Do the patterns of annotation frequency derived from InterProScan correspond to defining differences between the species? (2) Are the annotations that together form separate and clearly identifiable patterns for each species associated with functions that differ between species?

Given a number of samples, each belonging to one of *x* possible classes, a classifier attempts to learn the underlining sample-class mapping. Classes are labels with (in the present case) three possible values (*Panagrolaimus, Propanagrolaimus, Panagrellus*). Samples were n-dimensional vectors of annotation presence and absence. The classifier was trained on a set of samples, including an equal number of representatives from both classes and then tested on its ability to predict the class of an independent sample. We used 30 randomly drawn annotations from each representative species as dimensions in the classifier and ran 2000 iterations, each time selecting a different subset of training samples and test species. We tested variations relying on 50 or 100 dimensions (annotations), but found these to not change the overall pattern of results. Thirty dimensions were therefore chosen as the most efficient version.

Classification of Panagrolaimidae achieved 55 % accuracy (standard error of the mean [SEM] = 0.006; chance level classification = 50%) by a SVM based on randomly selecting 30 Pfam domains in each of the 2000 iterations through the database. The same approach yielded 58% mean accuracy (SEM = 0.006) classification for the Outgroup; the difference between classification performance for Panagrolaimidae and Outgroups was not statistically significant in a two-samples t-test (T(1999) < 1), showing that the classifier was not biased towards one class. More importantly, classification performance was significantly better than control classification (where sample-class mapping in the training set was permuted). Based on a permuted class-sample mapping in the training set, the classifier achieved a mean accuracy for Panagrolaimidae of 49 % (SEM = 0.006) and of 49% for the Outgroup (SEM = 0.008). The comparison between the real classifier incorporating true sample-to-class mappings in the training set and the permuted classifier was statistically significant for Panagrolaimidae: T = 7, p < 0.001, df = 1999; and for the outgroup: T= 7.4, p < 0.001, df = 1999 (see supplementary fig. 6).

During classification, we recorded annotations that were most often part of successful classification iterations. These lists were then intersected with results from the NHST analysis and interpreted with reference to identifying differences in the biology of each species group.

### Detection of Horizontal Gene Transfers

To detect candidate horizontal gene transfers (HGT), we used Alienness (Rancurel et al. 2017) to calculate an Alien Index (AI) as described in (Gladyshev et al. 2008; Flot et al. 2013). Briefly, all the Panagrolaimidae predicted proteins were compared against the NCBI’s nr protein database using BLASTp (Altschul et al. 1997) with an E-value threshold of 1e^−3^ and no SEG filtering. BLAST hits were parsed to retrieve associated taxonomic information, using the NCBI taxonomy database as a reference. For every Panagrolaimidae protein returning at least one hit in either a metazoan or non-metazoan species, we calculated the AI:

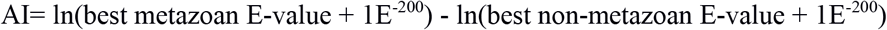

When either no metazoan or non-metazoan significant BLAST hit was found, a penalty E-value of 1 was automatically assigned as the best metazoan or non-metazoan E-value. To detect HGT events that took place in an ancestor of Panagrolaimidae or its close relatives, BLAST hits to Panagrolaimoidea (TaxID: 55746) and Aphelenchina (TaxID: 1182516) were skipped for the calculation of AI. No AI value could be calculated for proteins returning non-significant hits at all in NR. An AI > 0 indicates a better hit to a non-metazoan species than to a metazoan species and thus a possible acquisition via HGT. An AI > 30 corresponds to a difference of magnitude > 9.E^−14^ between the best non-metazoan and best metazoan E-values and is taken as strong indication of a HGT event (Rancurel et al. 2017).

All *Panagrolaimidae* proteins that returned an AI > 0 and that aligned with ≥70 % identity to a non-metazoan protein were considered as possible contaminants and were discarded from the analysis. Potential candidates were further validated by identifying gene models that contained spliceosomal introns, were surrounded by bona fide nematode genes, had mapped RNA-Seq reads (TPM estimates were calculated with RSEM (Li and Dewey 2011) incorporating bowtie2 (Ben Langmead and Salzberg 2012) mapping) and had codon usages similar to bulk genome values (supplementary fig. 7).

### Reconstruction of the timing of gene acquisitions via HGT

We used Mesquite v.3.01 (Maddison and Maddison) to reconstruct the timing of gene acquisition via HGT. By adding orthology information to HGT candidates, we built a matrix of presence/absence of the HGT candidates across the different *Panagrolaimidae* species. We mapped this matrix to the species phylogeny and included two additional outgroups (*Acrobeloides nanus* and *Bursaphelenchus xylophilus*). Based on presence/absence information, Mesquite then traced back ancestral presence / absence at each node, using a parsimony model. When ancestral presence/absence of a gene family at a given node was equally parsimonious, we arbitrarily considered the family as present. Indeed, it intuitively appears more likely to secondarily loose a gene that had been acquired by HGT ancestrally, than gaining it multiple times independently via HGT. HGT candidates that were species-specific were considered as acquired specifically in this species.

### Functional analysis of HGT candidates

Pfam annotations were retrieved for all the proteins putatively acquired via HGT, and domains that were conserved among the set of candidate HGT proteins of several Panagrolaimidae species identified (Supplementary Excel file HGT - PFAM). We also predicted putative functions basedon the presence of Pfam domains, using pfam2go (available at geneontology.org/external2go/pfam2go) and custom perl scripts. To make Gene Ontology annotations comparable, we mapped raw GO terms to the generic GOslim ontology. We used the GOSlimViewer, developed as part of AgBase (McCarthy et al. 2006), to map terms to the GOslim ontology (Supplementary Excel file HGT - GOSlim).

## Acknowledgements and Funding

The authors thank Isabel Goebel for generation time measurements, Janine Beyert and Andrea Kraemer-Eis for help in culturing nematodes. The authors are grateful to the GenoToul bioinformatics platform Toulouse Midi-Pyrenees for providing computing resources, as well as to Nordrhein-Westfalen for the use of the CHEOPS computing cluster at the University of Köln. This work was supported by the VolkswagenStiftung in their initiative for evolutionary biology in a grant to PHS, the DFG in a grant to TW under SPP1819 (supporting PHS), and by the European Research Council (ERC-2012-AdG 322790) in a grant to Max Telford (supporting PHS at UCL). AMB was funded by Science Foundation Ireland (Project RFP/EOB2506). EAMS thanks Dr. Ryan Gregory (University of Guelph) for providing access to his laboratory to conduct genome size estimations. She also thanks CONACyT for her Postdoctoral Fellowship (2010), and UAZ-2016-37096. CJC was funded under the Science Foundation Ireland (SFI) Stokes lecturer scheme (07/SK/B1236A) and the Biotechnology and Biological Sciences Research Council (BBSRC) Institute Strategic Programme Grant, Rumen Systems Biology, (BB/E/W/10964A01).

## Authors contributions

PHS: planned and conceived the study, managed the collaboration, analysed the data, supervised GS, wrote the paper

EGJD: analysed the HGT data, and planned and wrote the ms

AMB: analysed the anhydrobiosis and HGT data, conceived the study, supervised GO’M & BC, wrote the paper

A-MS: programmed the classifier, drew figures and contributed to writing

CJC: analysed the selection data, drew figures, and contributed to writing

SW: annotated the genomic data

ID: karyotyped species, drew figures, contributed to writing

GO’M: analysed anhydrobiosis data for *P. superbus*

BAC: conducted molecular experiments for transcriptomics

CR: computed and analysed the HGT data

GS: conducted genome size measurements

EAMS: conducted genome size measurements, wrote and revised the manuscript

UT: analysed raw sequencing data

MK: conceived parts of the study, co-supervised PHS

MAST: conceived parts of the study, co-initiated the collaboration, analysed the anhydrobiosis data

ES: conducted microscopic analyses, supervised PHS, and revised the manuscript

TW: analysed generation time data, wrote and revised the ms

MB: planned and conceived the study, analysed the data, and wrote the ms

